# gQuant: A Robust and Generalizable Algorithm for Identifying Normalizer Genes in qRT-PCR Data: A Case Study on Urinary Exosomal miRNAs

**DOI:** 10.1101/2023.11.25.568641

**Authors:** Abhay Kumar Pathak, Sukhad Kural, Shweta Singh, Lalit Kumar, Mahima Yadav, Manjari Gupta, Parimal Das, Garima Jain

**Author notes:** Correspondance –. Equal Contribution First Authors (Abhay Kumar Pathak, Sukhad Kural).

## Abstract

The emergent role of nucleic acid-based biomarkers, such as microRNAs (miRNAs), long non-coding RNAs (lncRNAs), and messenger RNAs (mRNAs), is becoming increasingly prominent in the realms of disease diagnostics and risk assessment. Quantitative reverse transcription PCR (qRT-PCR) is the primary analytical method for quantitative measurement of biomarkers. Yet, the relative infancy of non-coding RNAs’ (ncRNAs) recognition as biomarkers poses a challenge due to the absence of a consensus on a universally accepted normalizer gene, which is pivotal for accurate quantification. Current tools for selecting normalizer genes in qRT-PCR are fraught with limitations, including inadequate handling of null values, reliance on elementary statistical tools, use of a biased integrated approach, outlier sensitivity, and suboptimal graphical user interface for data visualization. These deficiencies underscore the necessity for a more nuanced and algorithmically balanced tool tailored to handle qRT-PCR datasets and facilitate the discernment of the most appropriate normalizer gene for specific datasets.

Addressing the identified challenges, we have developed ‘gQuant,’ a tool crafted to address the limitations present in existing methods. In ‘gQuant’ we employed voting classifiers as an ensemble technique that combines predictions from multiple statistical methods to make more accurate rankings than any individual statistical measures. The tool’s efficacy was substantiated through rigorous validation against datasets from the Gene Expression Omnibus (GEO) database and corroborated with experimental data derived from urinary exosomal miRNAs. Comparative analysis with existing tools revealed that their integrated methodologies could skew the ranking of normalizer genes, whereas ‘gQuant’ consistently yielded rankings characterized by lower standard deviation, reduced covariance, and enhanced kernel density estimation (KDE) values. Given ‘gQuant’s’ promising performance, normalizer gene identification will be greatly improved, improving the precision of gene expression quantification in a variety of research scenarios.

## 1. Introduction

Quantitative reverse transcription PCR (qRT-PCR) is a gold standard technique for the quantification of gene expression levels. Its importance has been magnified with the advent of microRNA (miRNA)-based biomarkers (Ye et al., 2019). The linchpin of comparative gene expression studies utilizing qRT-PCR is the judicious selection of normalizer genes for accurate data normalization (Smith et al., 2020). Although internal reference genes, such as ACTB, GAPDH, and 18s Ribosomal RNA (Suzuki et al., 2000), have been successfully employed at the tissue level for mRNA quantification, Researchers also opt to choose specific normalizers for individual experiments particularly where a universally accepted set of normalizers remains elusive such as in miRNA-based assays (Shen et al., 2011, Danese et al., 2017, Veryaskina et al., 2022). Although many miRNA-based studies used U6 as the most commonly used reference gene, it has variable expression and does not represent the optimal reference gene for miRNA analysis (Jacobsen et al., 2016).

Although, existing tools like RefFinder (Xie et al., 2023), use Ct values from experiments to determine the stability of genes. However such tools have limitations in addressing the unique challenges posed by the presence of null values, also it is based on a weighted approach that can be biased especially when dealing with complex biological variability. A detailed description of the limitations and approaches of currently available tools is provided in Table 1. Therefore, there is a need for robust, consistent, and generalizable algorithms for most stable gene identification.

**Table 1:** Comparative table for existing gene ranking tools advantages and limitations.

| Reference | Year | Tool Name | Advantages of Techniques | Limitations | Evaluation Matrix |
| --- | --- | --- | --- | --- | --- |
| (Vandesompele et al., 2002) | 2002 | GeNorm | It calculates a gene-stability measure M, for each gene and then derives a normalization factor based on the geometric mean of the selected best-performing reference genes.<br>The algorithm performs stepwise exclusion of the least stable gene, recalculating M values each time until the most stable genes are identified. | Statistical Robustness: It uses basic statistical metrics that might not encapsulate all the nuances in a complex dataset.<br>Assumption of Co-regulation: It assumes that the ideal reference genes are co-regulated, which might not be the case in all biological systems or conditions. | geometric mean, pairwise variation, Stepwise exclusion |
| (Andersen et al., 2004) | 2004 | NormFinder | Combined Estimation of Stable Gene Selection provides a generalized approach to understand data with different parameters | Sample Size must be above '8'<br>Candidate Size Must be above 5 | Intragroup and intergroup variation in gene expression levels |
| (Pfaffl et al., 2004) | 2004 | BestKeeper | Pairwise correlation covers inter-group validation, where other matrix measures are focused on intra-group validation | Data Assumption : it assumes that data is distributed normally<br>Outlier Sensitivity | GM, AM, SD, CV, MIN, Max and Pairwise Correlation |
| (Silver et al., 2006) | 2006 | $\Delta Ct$ | Intra-Group variation | Standard Deviation cannot do inter-group variation | Standard Deviation |
| (Li et al., 2015) | 2015 | GenEx Standard software | ACC SD (Accumulated Standard Deviation) captures both intra- and inter-group variability, potentially offering a more robust stability measure<br>Integration with NormFinder and GeNorm gives more robust results. | Initially assumed that data is normally distributed<br>Outlier Sensitivity | geNorm and NormFinder algorithms, Accumulated Standard Deviation |
| (Bruckert et al., 2016) | 2016 | Bio-rad CFX manager software | Focuses on both inter-group and intra-group variation | Metrics like Pairwise variation, geNorm and NormFinder are not integrated | Pairwise Variation,geNorm,NormFinder |
| (Marabita et al., 2016) | 2016 | summarized stability score (SSS) | Integrated approach which is able to understand the data carefully | NormFinder Struggles when the sample size is small, Outliers Sensitivity | GeNorm,NormFinder,Coefficient of Variation |
| (Hoja-Lukowicz et al., 2022) | 2022 | GenExpa | GenExpA introduces the use of a coherence score (CS) to validate the reliability of references.<br>GenExpA is a comprehensive tool that, based on the selected reference gene or pair of genes, calculates the relative target gene expression level, performs statistical analyses, and generates results | Assumes that data are preprocessed and well structured<br>It can not deal with missing values | NormFinder algorithm with a strategy of progressive removal of the least stable gene from the candidate genes in a given experimental model and in the set of daughter models assigned to it |
| (Xie et al., 2023) | 2023 | RefFinder | Integrated approach to find the Candidate reference gene | Weighted Approach is considered to be biased.<br>Assumed that data is normally distributed and ready to be processed<br>It does not have the functionality to deal with missing values | Various Tools like NormFinder, DeltaCt, BestKeeper, GeNorm |

MicroRNAs (miRNAs) have emerged as promising biomarkers for noninvasive diagnostic approaches in various cancers. Their stability, abundance, and presence in body fluids, such as urine, make them ideal candidates for cancer detection and monitoring (Condrat et al., 2020). Among these, urinary extracellular vesicles (uEVs) have garnered significant attention due to their cargo of stable miRNAs, reflecting the molecular signatures of their parent cells (Jahromi et al., 2023). One of such example is the exploration of uEVs to detect miRNAs released by tumors, especially in urological cancers like bladder cancer (BCa) and prostate cancer (PCa) in the search for non-invasive methods to detect and monitor the disease (Piao et al., 2021, Mugoni et al., 2022).

However, the absence of a robust and standardized normalizer led to the use of varied strategies for miRNA normalization, ranging from the use of spike-in controls (Sewer et al., 2014) like cel-miR-39 (Smith et al., 2020) to the employment of multiple endogenous miRNAs and their arithmetic means (Pagacz et al., 2020), as well as the pair ratio method (Lekchnov et al., 2018) or study specific normalizers (Jain et al., 2023, Veryaskina et al., 2022), raising questions about the comparability and reliability of results across studies.

Given this context, our effort to develop an efficient normalizer finding tool and its validation on uEV-miRNAs represents a timely and crucial endeavor. In this cross-disciplinary research initiative, we endeavor to synergize biological and mathematical fields to address this pressing issue. While searching for a standard normalizer gene for another study, we set to screen selected miRNAs based on existing literature. This initial assessment involved experimental qRT-PCR, followed by the use of existing tools like RefFinder. However, we encountered discrepancies in the results and limitations with these methods, which led us to develop a new and more robust analytical algorithmic model.

This paper introduces a novel and efficient algorithmic tool designed for analyzing qRT-PCR expression data to identify the most stable gene. We have validated our tool using a range of experimental data and existing datasets. Furthermore, we conducted a comparative analysis with current tools, pointing out their limitations, and detailed the methodological thoroughness that went into the development of this innovative analytical approach.

## 2. Material and Method

### 2.1. Platform for Mathematical Analysis

We have used the Jupyter Notebook interface to write scripts in ‘python’, which is a web-based interactive computing environment that allows the execution of live code, embedding of visuals, and explanatory text all in one document. Libraries used were, Pandas: Used for data manipulation and analysis, NumPy: A fundamental library for numerical computations in Python, SciPy: Used for scientific and technical computing, Scikit-learn: This library is used for machine learning and data mining tasks.

### 2.2. Subjects

During the period spanning October 2022 to August 2023, urine specimens were taken from individuals who were diagnosed with BCa, PCa and asymptomatic healthy control subjects were recruited from Department of Urology, Institute of Medical Sciences, Banaras Hindu University, India. Controls were chosen to closely match patients in age (range 45-60) and absence of prior cancer history. Consent for participation was obtained from all the individuals involved. The study began post securing required ethical approvals from relevant authorities (Institute of Science, BHU, Varanasi & Institute of Medical Sciences, BHU, Varanasi).

### 2.3 Exosomal miRNA Isolation

Norgen Urine Exosome RNA Isolation Kit (Catalog #47200, Norgen Biotek Corp.) was used to extract uEV-miRNAs. We chose a 10 mL urine sample as the starting volume, following the kit’s instructions, and processed the sample. RNA quality and concentration were then determined using NanoDrop (Thermo Scientific, USA).

### 2.4 cDNA synthesis

Reverse transcription of RNA samples to cDNA was performed using RevertAid First Strand cDNA Synthesis Kit (Catalog #K1622, Thermoscientific, USA). We used miRNA-specific stem-loop primers to reverse transcribe targeted miRNA only. Primer details are provided in Supplementary Table 1

### 2.5 Quantitative reverse transcription PCR

We conducted qRT-PCR using the Maxima SYBR Green/ROX qPCR Master Mix (2X) (Catalog #K0221, Thermo Scientific, USA). The reaction was performed using Applied Biosystems QuantStudio 6 Flex Real-Time PCR System.

## 3. Process and Development of gQuant

### 3.1. Current tool Limitation

I. Missing Value: The lack of methods to handle missing values in existing tools, such as Gnorm, DeltaCt, BestKeeper, RefFinder, and GenExpa, raises concerns about the final ranking produced by these tools. We strengthened this weakness by including an extra preprocessing unit in our tool to reduce the dangers associated with working with NULL values.
II. Different approaches: Current tools use diverse statistical matrices that provide different understandings of data, its nature, and interpretation. Whereas, our tool uses four evaluation matrices : Kernel Density Estimation, Standard Deviation, Covariance, and Geometric Mean, to combine the diversity of observations and enable multidimensional analysis from a distinctive angle.
III. Scaling: On any integration system, using literal values of different matrices could lead to dominance, as it is in existing tools. In democratic voting, we have employed standard scaling to standardize the results into the interval [0,1], preventing the dominance of a single matrix. It will guarantee that none of the four matrices can affect the voting process.
IV. Interactive Graphics: During the gene ranking procedure, visual depictions serve as a critical tool for the users. They allow us to understand the core characteristics of the data, its fluctuations, its distribution, and the density of its scatter points. We employed a box plot that displays the expression of various genes over distinct intervals, indicating the data scope and distribution. Furthermore, we incorporated a KDE (Kernel Density Estimation) plot to illustrate the density of data points (and their specific numerical values) as well as their spread range.
V. Democratic Voting-Based Integration: Traditional tools such as RefFinder have relied on weighted schemes that could introduce biases by assigning predetermined weights to rank genes. To address this issue, our tool adopts a democratic voting strategy, where each gene competes for votes based on its characteristics. The gene receiving the majority of votes is selected, and an iterative ranking index is constructed. This method ensures a fair and balanced approach to gene ranking, minimizing potential biases inherent in weighted systems.

### 3.2 Tool Development

The algorithm aims to address the challenge of gene ranking using a multi-metric approach that combines four different statistical measures to iteratively rank gene’s stability. This algorithm utilizes Standard Deviation, Geometric Mean, Covariance, and Kernel Density Estimation to select and rank genes iteratively from a high-dimensional dataset.

#### I. Data Preprocessing

During the initial preprocessing stage, as the input dataset is introduced, the program embarks on data preprocessing by identifying and quantifying the missing values or null values within each gene column. Subsequently, it calculates the ratio of available values to missing values (denoted as NA) for every individual gene column. This ratio is instrumental in determining the subsequent action: should the ratio transcend a user-specified threshold,the tool engages in a missing value imputation process. Here, it substitutes the missing value (NA) with the median of the available values within the respective gene column. Conversely, if the ratio is found to be beneath the stipulated threshold, the program autonomously eliminates the particular column from the dataset, a mechanism applied iteratively across all gene columns within the dataset. This imputation strategy is predicated on an assumption of non-normal data distribution within the miRNA dataset. Ratio of available values to missing values is problem specific,and could be chosen manually. For our result and validation we have taken the 8:1 ratio as a threshold for preprocessing. This detailed procedure represents the inaugural part of the tool’s operational framework, formulated in a scientific discourse as Part A (Figure 1).

**Figure 1:**
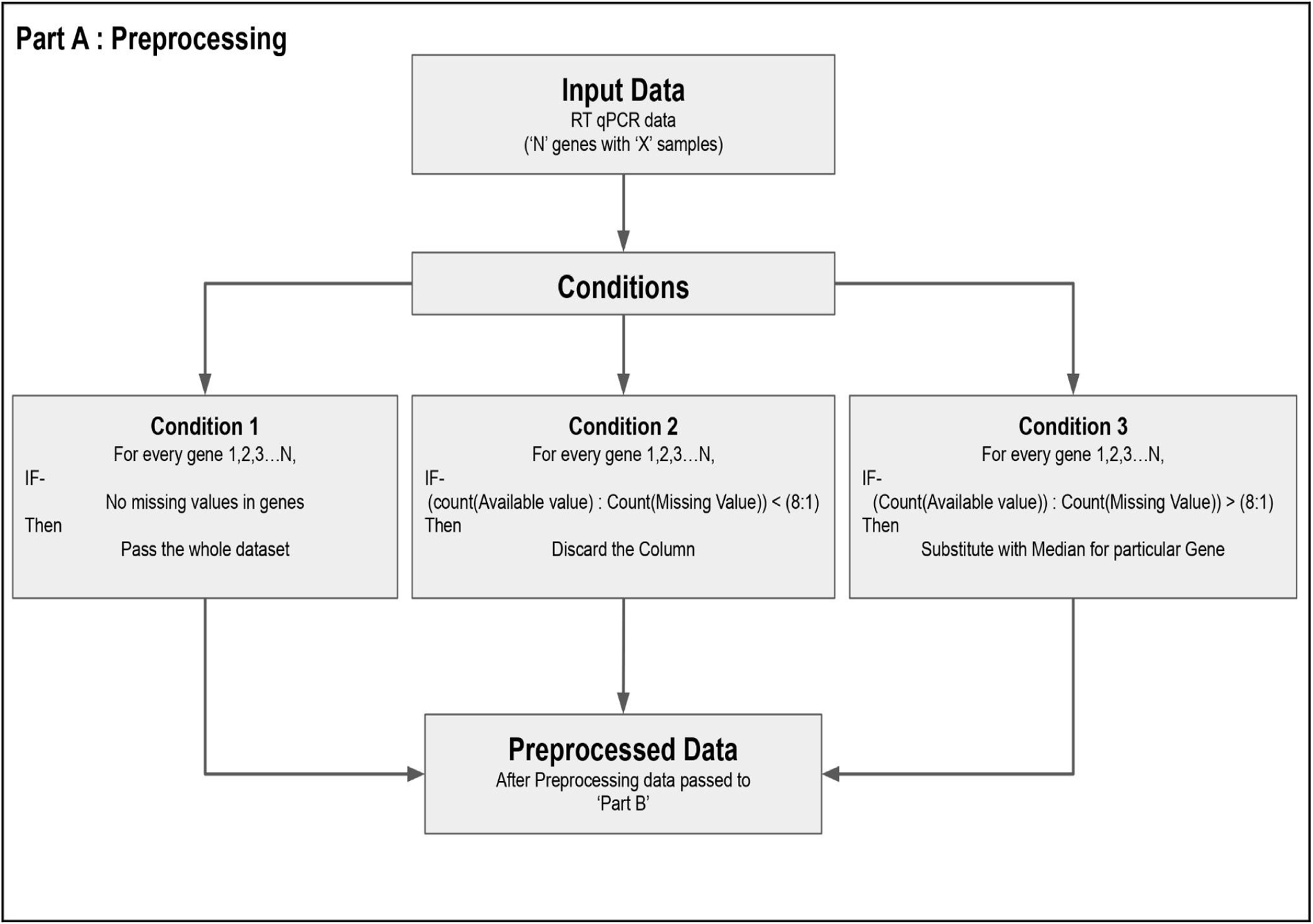
The workflow of Part A : Preprocessing.

### II. Selection of the most stable gene

In an effort to develop a model that synergistically combines the strengths of existing techniques while mitigating their limitations, optimizing the performance evaluation matrix for more nuanced and robust gene selection, we followed a systemic approach. As explained in the following sections.

**Figure 2:**
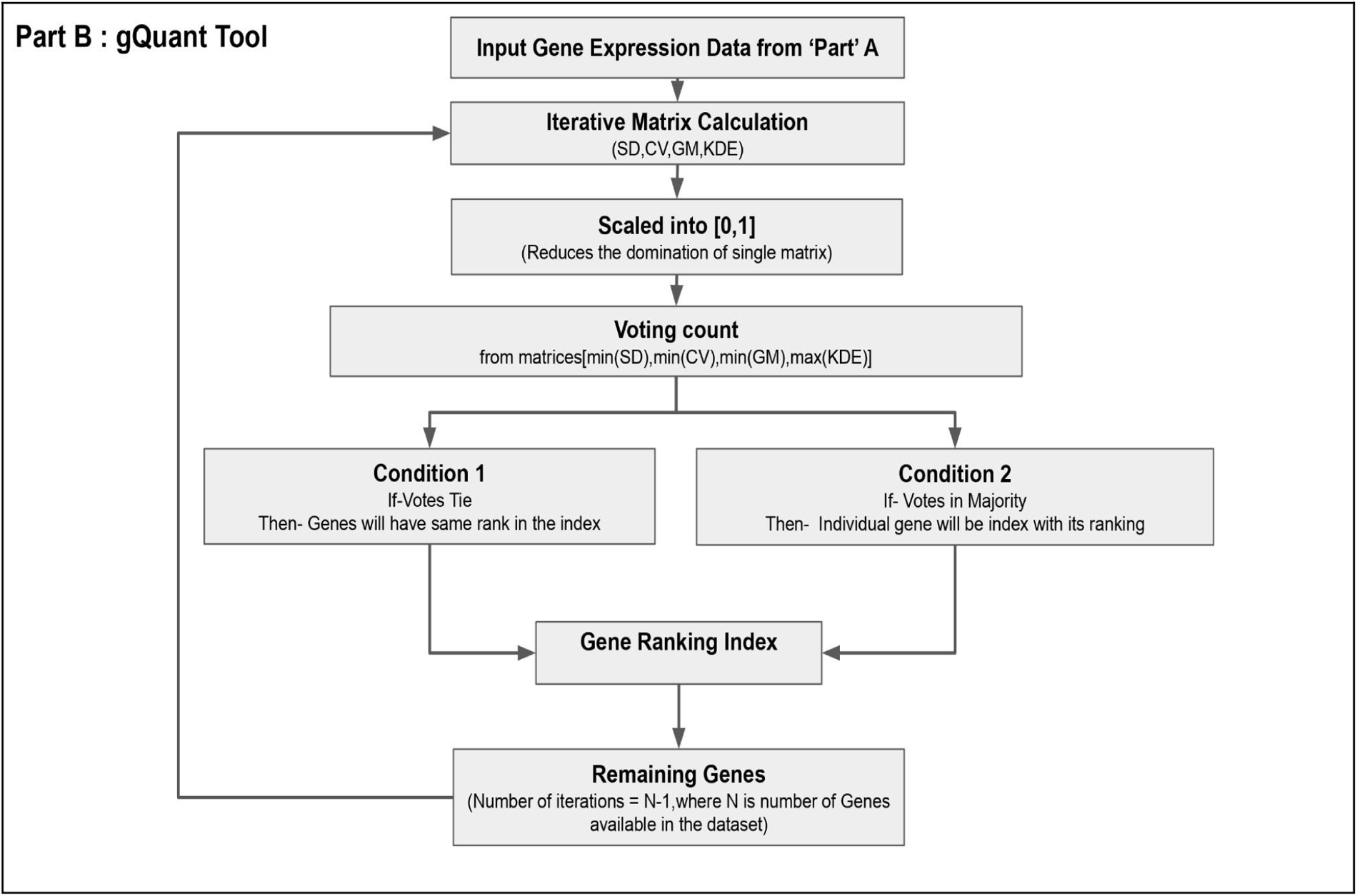
gQuant Tool workflow ‘Part B’ for the processing of preprocessed data.

The tool first uses “Part A’s” preprocessed data as input (Figure 2). It then computes all the matrices, scaling the results from the four matrices into a range of [0,1] to prevent any disproportionate impact of individual matrices on the voting outcome. Two things could happen in order for us to use a majority voting integration strategy to determine which gene is the most stable across the dataset: When there is a tie in the first round of voting, the tied genes are given the same ranking in the index; if there is a majority vote for a particular gene, it is saved on the ranking index and the remaining genes repeat the process until only one gene is left.

Metrics Used in Tool:

I. Standard Deviation (SD): In our model for the selection of the most stable gene, standard deviation was employed as a key metric. We used SD to quantify the level of gene expression variability across various experimental groups. The algorithm computes the standard deviation within groups for each gene’s expression levels, which serves as an initial filter in identifying potentially stable genes. Low standard deviation shows that the gene’s expression levels are tightly clustered around the mean, conversely, genes with high standard deviation are loosely clustered means they are unstable for being chosen to be a reference gene. The standard deviation *SD*(*g_i_*) of a gene *g_i_* with *m* samples is given by:

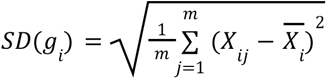

where *X_ij_* is the expression level of gene *g_i_* in sample *j*, and 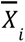 is the mean expression of *g_i_*
II. Geometric Mean (GM): The geometric mean is a measure of central tendency that is more robust to extreme values (outliers) compared to the arithmetic mean.GM is used to identify genes that maintain a consistent expression level across all samples, since a low GM would imply that at least one sample has a low expression level. The geometric mean *GM*(*g_i_*) for a gene *g_i_* is

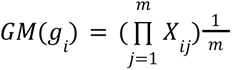

Where *m* is the number of samples.
III. Covariance (CV): Covariance provides a measure of the degree to which two variables change together. The Covariance Mean is an average of the covariances between one gene and all other genes. High covariance mean values indicate genes that are likely to be involved in similar biological quantification, and therefore may be of particular interest. The Covariance *CV*(*g_i_*) for a gene *g_i_* is

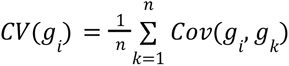

Where *Cov*(*g_i_,g_k_*)is the covariance matrix between *g_i_* and *g_k_*
IV. Kernel Density Estimation (KDE): Kernel Density Estimation is a non-parametric way to estimate the probability density function of a random variable. In the context of gene ranking, it is used to estimate the distribution of expression levels for each gene.KDE serves to identify genes that have expression levels concentrated around the mean, which may be of biological significance. The Kernel Density Estimation *KDE*(*g_i_*) for a gene µ*_i_* is

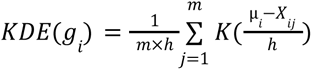

Where *K* is the Kernel function, *h* is the bandwidth, and µ*_i_* is the mean of *g_i_*

Mathematical formulation of gQuant tool algorithm:

Mathematical Notation:

● *G*: Set of Genes *g*_1_, *g*_2_, *g*_3_……,*g*_n_
● *X*: Matrix of gene expression values, where *X_ij_* denotes the expression value of gene *g_i_* in Sample *j*.
● *S*: Standard deviation vector *S = [S_1_, S_2_, S_3_*………,*S*_n_]
● *GM*: Geometric mean vector *GM = [GM_1_, GM_2_, GM_3_*………,*GM*_n_]
● *COV*: Covariance mean vector *COV= [COV_1_, COV_2_, COV_3_*………,*COV*_n_]
● *KDE*: Kernel density estimation vector *KDE = [KDE_1_, KDE_2_, KDE_3_*………,*KDE*_n_]
● *R*: Ranking index list 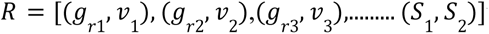 where *g_ri_* is a gene and *v_i_* is gene ranking

Initialization: Read the gene expression data, extracting names and numerical values for further processing.

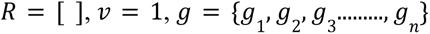

Scaling: All metrics are scaled to ensure that no single metric dominates the ranking procedure.

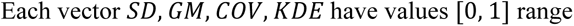

Iterative Ranking Metric Calculation: For each remaining gene *i* = 1 to *n* − 1, all four metrics are calculated.

Voting: Each metric elects the gene with the “least” value (except for Kernel Density, which looks for the maximum).

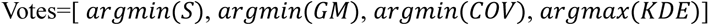

Majority Rule: The gene that appears the most frequently in these elections is removed from the data and ranked.

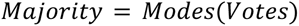

Update *R*[ ] majority with *v*, then remove majority from *G*, with increment of *v*

Tie Breaking: If multiple genes tie for the most votes, all are removed and given the same rank.

Recursion: Steps are repeated until only one gene remains.

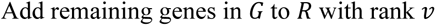

## 4. Overview of Validation Dataset

To ascertain gQuant’s precision, three distinct qRT-PCR datasets were utilized:

I. Dataset One: Derived from research on OvCar-3 and PC-3 cancer cell lines, this dataset encompasses transcriptomic profiling of 84 genes pertinent to cell regulation and five conventional housekeeping genes. The data, accessible via GSE57888, serves as a benchmark for mRNA-based qRT-PCR analysis, offering validation for gQuant through non-normalized data.(Tilli et al., 2014)
II. Dataset Two: Dataset Two: We chose a dataset GSE239868 with more detailed variables, such as the expression of 1,066 human miRNAs and n=36 tracheal aspirate samples, for a comparative efficiency assessment of gQuant. This database included data for a comparative analysis on the management of missing values and allowed us to test the appropriateness of our method on miRNA background. This dataset’s unnormalized Ct values provide a strict testing environment (Silveyra et al., 2023).
III. Dataset Three: In an endeavor to construct a more robust analytical framework, we decided to explore the potential of microRNA (miRNA)-based normalizers as an alternative to the well-established messenger RNA (mRNA)-based normalizers. Given the nascent state of miRNA-based normalization data, we focused specifically on uEVs. Sample collection and processing are explained in the method section. The miRNAs set selected for study validation underwent a thorough selection process, combining a comprehensive literature review and the Qiagen’s Human Urine Exosomes Focus miRCURY LNA Panel, and was further supported by analysis of RNA sequencing data. This investigation led to the selection of miR-16-5p, miR-10b-5p, 30b-5p, and miR-30d-5p (Bryzgunova et al., 2016) (Rodríguez et al., 2017) (Fredsøe et al., 2019). Additionally, upon evaluating the expression levels of the Let7c cluster genes from another ongoing study, we observed high stability for let-7c-5p across both diseased and control samples, leading to its inclusion in subsequent investigations. Therefore, Our preliminary study proceeded to focus on the selected miRNAs: let-7c-5p, miR-16-5p, miR-10b-5p, miR-30a-5p, and miR-30d-5p. Using qRT-PCR, we assessed their expression in uEVs samples of BCa (n=9), PCa (n=6), and control samples (n=3). To evaluate the stability of miRNA expression both with the publicly available tools RefFinder and using gQuant (Table 5). Among the evaluated miRNAs, let-7c-5p has emerged as the most promising candidate for normalizer functions. Subsequently, expression levels for let-7c-5p miRNAs in uEVs were quantified in an expanded sample set (n=30) using qRT-PCR (Figure 5). Rigorous quality control measures, including primer efficiency and melt curve analysis, were implemented, and any data with Ct values over 38 were excluded. The stability of the miRNA expression was validated using both RefFinder and gQuant, solidifying let-7c-5p’s potential as a reliable normalizer.

## 5. Result and Validation: Performance of gQuant in Control Gene Identification

### 5.1. Baseline Evaluation with Established Endogenous Controls

We employed an external qRT-PCR dataset from the Gene Expression Omnibus (GEO) to extensively assess the performance of our model. A dataset (Dataset One) comprising the expression values of reputable endogenous controls was employed for this assessment. Table 2 presents the results, where the normalizer that our model identified has the greatest ranking is shown. Traditional mRNA normalizers consistently ranked in the top 5 in our analysis. We were able to verify the model’s applicability in a range of experimental situations and assess the model’s performance on independent data.

**Table 2:** Ranking index of most stable gene obtained using gQuant on dataset One(GSE57888).

| Ranking Index | Genes |
| --- | --- |
| 1 | HPRT1 |
| 2 | B2M |
| 3 | RPL13A |
| 4 | ACTB |
| 5 | GAPDH |
| 6 | TWIST1 |
| 7 | PLAU, VEGFA, MDM2,ANGPT2 |
| 8 | MTSS1,PPC3,TIMP3,HGDB |
| 9 | THBS1 |
| 10 | CDKN2A |
| 11 | CDKN2A.1 |

### 5.2. Assessing gQuant’s Stability with Heterogeneous miRNA Expression

A Comparison with RefFinder – To assess gQuant’s sensitivity to expression variability and the presence of outliers and missing values, we conducted a comprehensive sensitivity analysis using a subset of dataset Two. The subset included 51 genes out of 1066 genes for all 36 sample sizes. The dataset included C. elegans miR-39 as an exogenous spiked-in control, six snoRNA/snRNA genes, and a Positive PCR Control (PPC) for PCR performance benchmarking. The comparative analysis is detailed in Table 3.

**Table 3:** Comparison of ranking obtained by gQuant and RefFinder using dataset Two (GSE239868).

| Gene | gQuant Ranking | RefFinder Ranking |
| --- | --- | --- |
| cel-miR-039 | 6 | - |
| cel-miR-39.1 | 11 | 17 |
| SNORD61 | 8 | 4 |
| SNORD68 | 9 | 17 |
| SNORD72 | - | 40 |
| SNORD95 | 12 | 8 |
| SNORD96A | 10 | 22 |
| RNU6-2 | 10 | 27 |
| miRTC | 6 | 3 |
| PPC.1 | 1 | - |
| PPC | 2 | 1 |

which showcases the ranking of normalized genes identified by the model. gQuant’s performance enhancement is evidenced by the superior ranking of normalization genes, indicating its robustness in handling data anomalies, including outliers and missing values. Additionally, the lower ranking of the spiked-in control C-miR-39 underscores the potential impact of manual errors on data normalization processes. ‘SNORD 72’ transcends the specified threshold ratio of 8:1,so the rank is undermined.In RefFinder column, the miRNA ‘cel-miR-039’s Rank was undetermined.

### 5.3 Selection and Validation of let-7c-5p as the Superior Normalizer Using Dataset Three

Utilizing Dataset Three, which focused on the expression profiles of miRNAs including let-7c-5p, miR-16-5p, miR-10b-5p, miR-30a-5p, and miR-30d-5p, our goal was to determine the most reliable normalizer among the candidates. gQuant accorded the highest stability ranking to let-7c-5p, as presented in Table 4.

**Table 4:** Ranking index for most stable gene predicted using gQuant on Dataset Three.

| Ranking Index | Name |
| --- | --- |
| 1 | Ct Mean (let-7c-5p) |
| 2 | miR-30a-5p |
| 3 | miR-10b-5p |
| 4 | miR-30d-5p |
| 5 | miR-16-5p |

The validity of this ranking was reinforced through evaluative measures of distribution (Figure 3).

**Figure 3:**
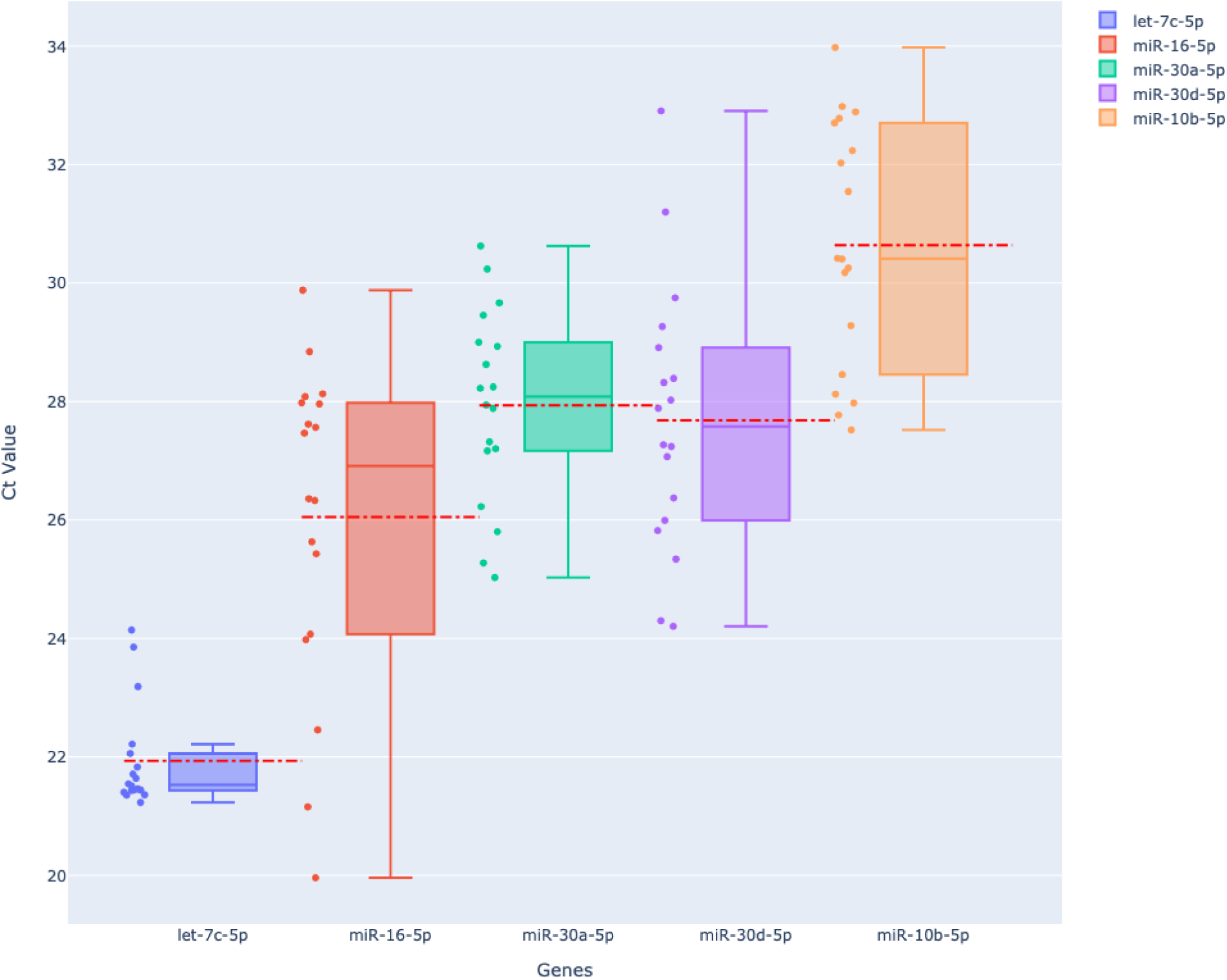
Ct value distribution for genes, depicted as a box plot with a strip plot overlay. Cycle threshold (Ct) values for miRNA expression ‘Ct Mean (let-7c-5p)’, ‘miR-16-5p’, ‘miR-30a-5p’, ‘miR-30d-5p’, and ‘miR-10b-5p’ are distributed as shown in this figure. Genes are plotted on the x-axis, and their corresponding Ct values are quantified on the y-axis. The interquartile range (depicted by boxplot), median (depicted by colored line in boxplot),mean(depicted by red dashed line) and outliers are displayed in each box plot, and the strip plot offers more information on the distribution of the data.

The gene “let-7c-5p” served as a reference, with its tighter distribution and lower value, showing a more consistent expression level. Conversely ‘miR-30d-5p’ and ‘miR-10b-5p’ demonstrate a broad range of expression levels with several points as outliers. The data points overlay on the boxplot allows for the detailed analysis of the spread and particular variation within different genes. And the concentration of data points via Kernel Density Estimate (KDE) in figure 4.

**Figure 4:**
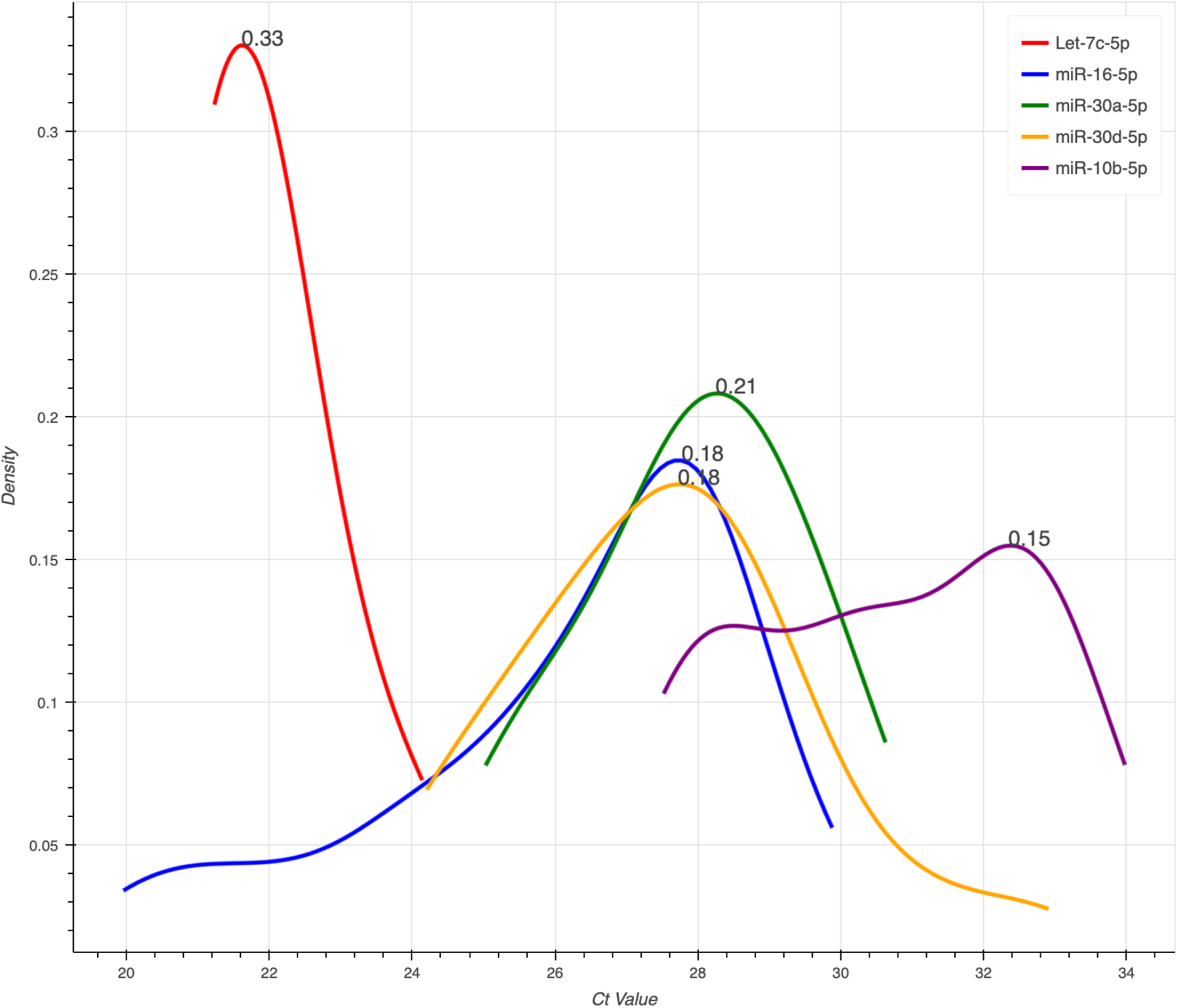
KDE Plot of Ct Values Across Genes. A Kernel Density Estimation (KDE) plot of cycle threshold (Ct) values for the genes “Ct Mean (let-7c-5p)”, “miR-16-5p”, “miR-30a-5p”, “miR-30d-5p”, and “miR-10b-5p”. The estimated density is displayed on the y-axis, Ct values are displayed on the x-axis. The map also shows different peaks and distribution patterns indicating different features of their expression.

KDE plot shows the distribution of Ct Values of different genes from our dataset.The X-axis represents the Ct values which are quantifying characteristics of gene expression level and Y-axis represents the estimated density. Depicts the distribution of Ct values for various Genes labeled as “Ct Mean “let-7c-5p”, “miR-16-5p”, “miR-30a-5p” “miR-30d-5p”, and “miR-10b-5p”.The Red curve shows the averaged Ct values for let-7c-5p with KDE value of 0.33,displaying a prominent peak at approximately 22, showing a mode of expression level for this gene. For other genes they show different distribution patterns and peaks, indicating different expression nature.

We used a Gaussian kernel with calculated bandwidth to correctly reflect the distribution. KDE highlights the major aspects like dispersion,multimodel Ct Values and central tendencies to get insights and expressive nature of the individual genes. Further validation was pursued through an expanded study encompassing a diverse set of uEVs samples, including PCa (n=22), BCa (n=30), BPH (Benign Prostatic Hyperplasia) (n=7), and Control (n=16). An analysis of let-7c-5p expression across these biological cohorts revealed minimal variability, thereby substantiating its consistent expression irrespective of the biological context.

In boxplot(figure 5), illustrating the distribution of Cycle threshold(Ct) across four different classes BPH, BCa, Control and PCa where we have compared control with BCa and BPH with PCa.To see the difference, we formulated a NULL hypothesis “that there are no significant differences between control with BCa and BPH with PCa”. As we can see by doing the t-test, we got p-values of 0.1542 for control with BCa comparison and a p-value of 0.1688, which are not significant. So we can say there are no statistical differences between both comparisons.

**Figure 5:**
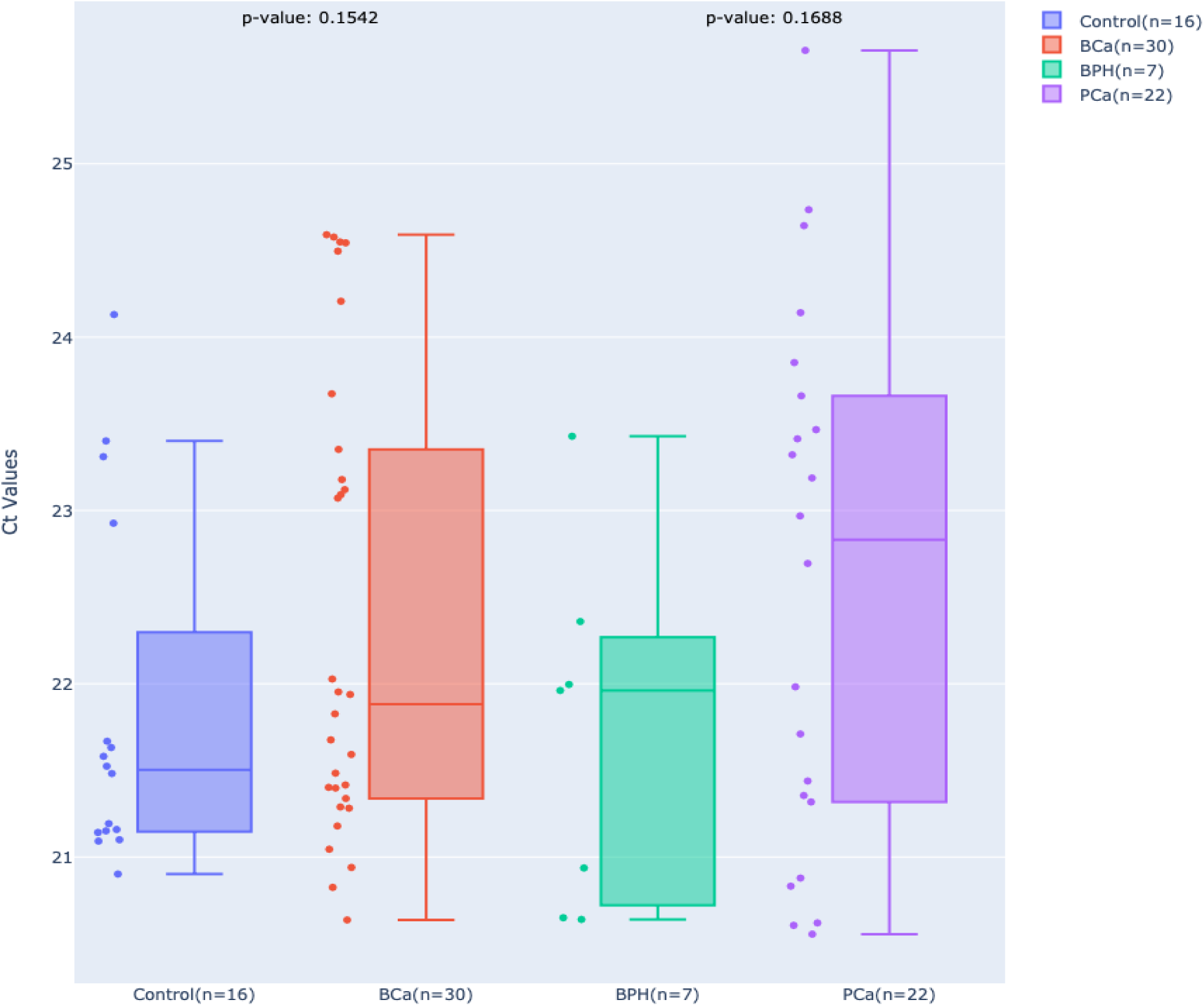
Distribution of Cycle threshold(Ct) across four different classes BPH, BCa, control, and PCa where we have compared ‘control with BCa’ and ‘BPH with PCa’.

### 5.4 Compared with refFInder and other existing tools available for gene ranking

Here we have attached figure 6, a snapshot of RefFinder Tool result on dataset two, showing ‘miR-30a-5p’, as most stable genes followed by ‘miR-10b-5p’, ‘let-7c-5p’, ‘miR-30d-5p’, ‘miR-16-5p’. Which is different then gQuant Tool ranking.

**Figure 6:**
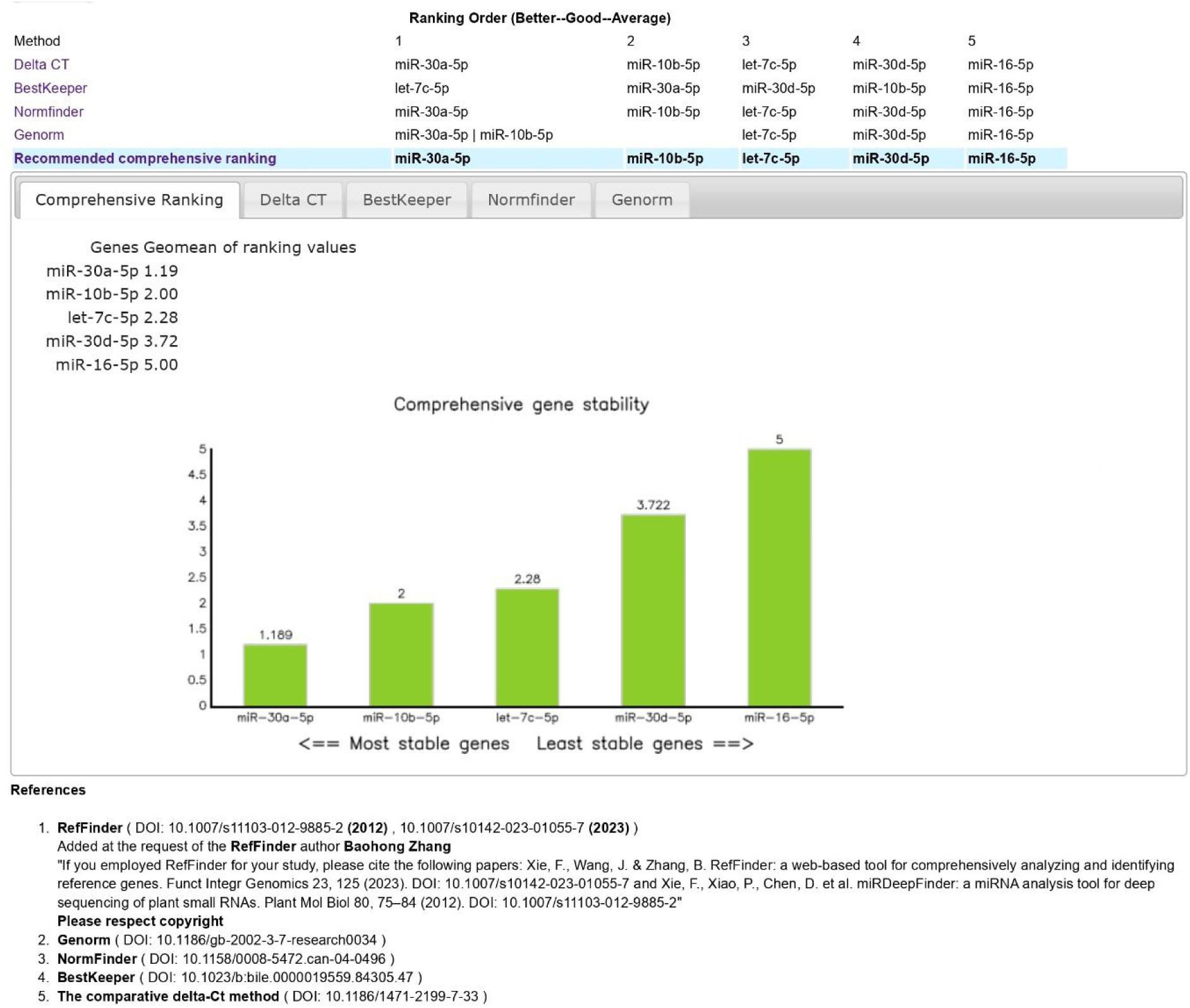
RefFinder ranking index on dataset three (Rank followed as miR-30a-5p, miR-10b-5p, let-7c-5p, miR-16-5p)

On dataset two, we have presented a stable gene ranking in table 5, according to gQuant and other tools available. The compact distribution of let-7c-5p values (Figure 3), high density of data points (Figure 4), and east variation among biological data points (Figure 5), validate the stability of let-7c-5p as predicted by gQuant. However, the ranking index given by RefFinder and other tools does not follow the same pattern.

**Table 5:** Gene ranking index of dataset three Using gQuant,RefFinder, Delta Ct, BestKeeper, NormFinder and GeNorm.

| Ranking Index | gQuant | RefFinder | Delta Ct | BestKeeper | NormFinder | GeNorm |
| --- | --- | --- | --- | --- | --- | --- |
| 1 | **let-7c-5p** | miR-30a-5p | miR-30a-5p | let-7c-5p | miR-30a-5p | miR-10b-5p |
| 2 | **miR-30a-5p** | miR-10b-5p | miR-10b-5p | miR-30a-5p | miR-10b-5p | miR-30a-5p |
| 3 | **miR-10b-5p** | let-7c-5p | let-7c-5p | miR-30d-5p | let-7c-5p | let-7c-5p |
| 4 | **miR-30d-5p** | miR-30d-5p | miR-30d-5p | miR-10b-5p | miR-30d-5p | miR-30d-5p |
| 5 | **miR-16-5p** | miR-16-5p | miR-16-5p | miR-16-5p | miR-16-5p | miR-16-5p |

## 6. Discussion

In this work, we present a novel algorithm called ‘gQuant’ that aims to enhance the methodology of stable reference gene identification in gene expression investigations utilizing the widely used laboratory method of qRT-PCR. “gQuant” is a powerful and versatile algorithmic tool designed to overcome the drawbacks of current approaches. The tool employs additional pre-processing of data to ensure efficient dealing of missing values.

Our approach leverages democratic voting classifiers, combining predictions from multiple statistical methods to yield more accurate rankings than any individual measure. This technique presents a graphical demonstration to depict data fluctuation distribution and density.

Our model underwent rigorous validation using external qRT-PCR datasets from the Gene Expression Omnibus (GEO) and in-house generated datasets. Dataset One represents traditional qPCR data consisting of amplification values for target genes and conventional normalizer genes. when we ran this data through gQuant, as expected all the conventional normalizer genes scored top five ranks. This validation not only approves the quality of the gQuant but also shows its applicability on an independent dataset. dataset two represents a larger data set with a high number of target genes and samples. Since this dataset has null values in each gene column, the validation of this dataset reflected gQuant’s ability to handle null values as compared to Reffinder. The list of conventional normalizers used in this dataset includes the spike in controls, the family of SNORD (-61, -68, -72, -95, -96A), U6 small nuclear RNA, one unknown gene depicted as miRTC, and PCR positive control. When a subset of dataset Three, was run through gQuant, above mentioned normalizers could score ranks between 1 to 12. Whereas Reffinder’s rank indices went as low as 27. In both the cases, however, SNORD72 could not be ranked. gQuant eliminated SNORD72 as its null-value ratio exceeded 8:1 whereas the reason for Reffinder’s undetermined result for SNORD72 is not known. Nevertheless, we could see variations among the results obtained from Reffinder and gQuant and improved ranking indices of conventional normalizer genes approves the better efficiency of gQuant.

Dataset three represents amplification data from uEV-miRNAs, where there is the absence of any classical normalizer genes or small nuclear RNAs. To generate this dataset, we isolated uEV-miRNAs and collected qRT-PCR data for 5 selected miRNAs. These 5 miRNAs were previously used as stabilizers in uEV-based studies. Amplification data for these 5 miRNAs were first collected using a small cohort. This generated data was then run through gQuant, which distinctly identified let-7c-5p as normalizer. Whereas RefFinder placed let-7c-5p on rank 3 when the Same dataset was checked. To check the precision of this result, we generated let-7c-5p expression data on a larger cohort with 4 biological groups. The high p-value among the biological group supports the fact that the expression of let-7c-5p shows low variance among the biological groups. Apart from this, high-density points as depicted by the KDE curve and compact distribution of let-7c-5p as compared to other genes, also support the fact that let-7c-5p should be ranked highly stable gene among the given dataset.

These validations demonstrate our tool’s robustness. Nevertheless, gQuant is not without its restrictions. For example, if the criteria for missing data are excessively stringent, it may miss possible reference genes. At this time, multiplex-RT-PCR and RNA sequencing data are not supported by gQuant’s validation; it is exclusively verified for qRT-PCR data. It should be the goal of future research to apply it to more data kinds.

All things considered, gQuant provides a more accurate method for locating stable reference genes in gene expression research. This progress is essential for fields such as molecular biology research and disease diagnosis. With its notable advancement in gene expression analysis, gQuant has great promise for expanding our knowledge in the realms of molecular and medical research.

## Author Contribution

Research concept and design: GJ; Coding of gQuant: AP; Patient Sample Collection: SK,SS and LK; Pathological analysis MY; Collection and/or assembly of data: SK, AP, SS; Data analysis and interpretation: GJ, AP; Writing the article: GJ, SK, AP; Critical revision of the article: GJ, MG and PD. All the authors have approved the submitted version and agree to be personally accountable for the author’s own contributions and for ensuring that questions related to the accuracy or integrity of any part of the work.

