## Supplementary Table 1 for "gQuant: A Robust and Generalizable Algorithm for Identifying Normalizer Genes in qRT-PCR Data: A Case Study on Urinary Exosomal miRNAs"

| Gene Names | Primer Names | Sequences | Length/nt |
|---|---|---|---|
| let-7c-5p | S-L Primer[a] | GTCGTATCCAGTGCAGGGTCCGAGGTATTCGCACTGGATACGACAACCAT | 50 |
| | F-Primer[b] | AACACGCTGAGGTAGTAGGTT | 21 |
| miR-16-5p | S-L Primer | GTCGTATCCAGTGCAGGGTCCGAGGTATTCGCACTGGATACGACCGCCAA | 50 |
| | F-Primer | AGGGCGTAGCAGCACGTA | 18 |
| miR-30a-5p | S-L Primer | GTCGTATCCAGTGCAGGGTCCGAGGTATTCGCACTGGATACGACCTTCCA | 50 |
| | F-Primer | AACGGCTGTAAACATCCTCG | 20 |
| miR-30d-5p | S-L Primer | GTCGTATCCAGTGCAGGGTCCGAGGTATTCGCACTGGATACGACCTTCCA | 50 |
| | F-Primer | AACACGCTGTAAACATCCCC | 20 |
| miR-10b-5p | S-L Primer | GTCGTATCCAGTGCAGGGTCCGAGGTATTCGCACTGGATACGACCACAAA | 50 |
| | F-Primer | AACACGCTACCCTGTAGAACC | 21 |
| | Uni-R Primer[c] | CCAGTGCAGGGTCCGAGGTA | 20 |

a: Stem-Loop primer b: Forward primer c: Universal-Reverse PCR Primer
